# Stimulus presentation rates affect performance but not the acquired knowledge – Evidence from procedural learning

**DOI:** 10.1101/650598

**Authors:** Mariann Kiss, Dezso Nemeth, Karolina Janacsek

**Affiliations:** Department of Cognitive Science, Budapest University of Technology and Economics, Budapest, Hungary; Institute of Psychology, ELTE Eötvös Loránd University, Budapest, Hungary; Brain, Memory and Language Research Group, Institute of Cognitive Neuroscience and Psychology, Research Centre for Natural Sciences, Hungarian Academy of Sciences, Budapest, Hungary; Lyon Neuroscience Research Center (CRNL), Université Claude Bernard Lyon 1, Lyon, France

**Keywords:** implicit learning, probabilistic learning, stimulus presentation rate, response-to-stimulus interval (RSI), competence vs. performance

## Abstract

Presentation rates – the tempo in which we encounter subsequent items – can alter both our behavioral and neural responses in cognitive domains such as learning, memory, decision-making, perception and language. However, it is still unclear to what extent presentation rates affect the momentary performance versus the underlying cognitive function or mental representation. Here we systemically tested the effect of presentation rate on performance versus competence in procedural learning – a fundamental cognitive function that underlies the acquisition of skills and habits. Procedural learning was assessed by a probabilistic sequence learning task, in which learning typically occurs implicitly. Stimulus presentation rates were systematically manipulated by the Response-to-Stimulus-Intervals (RSIs). Participants were assigned to a fast (120ms) RSI or a slow (850ms) RSI group during the Learning phase, and were tested with both RSIs 24 hours later (Testing phase). We also tested whether they gained explicit knowledge about the sequence or their knowledge remained implicit. We found that the slower RSI led to weaker performance in the Learning phase. Importantly, however, the results of the Testing phase showed that this effect was primarily due to the expression of the acquired knowledge and not the learning itself. Thus, the slower presentation rate affected performance rather than competence. The acquired knowledge remained implicit in both groups, regardless of the presentation rate during learning. These findings call for tasks that can dissociate competence from performance both in experimental and clinical settings. Additionally, our findings can help design more efficient training protocols in education and clinical rehabilitation.

## 1 Introduction

In multiple domains of cognition, such as learning, memory, decision-making, perception and language, the presentation rate – that is, the tempo in which we encounter subsequent items (events) – is critical from both a theoretical and a methodological perspective. Theoretically, presentation rates can determine how our mind processes those items, discovers potential relationships among them and even binds them together (Destrebecqz & Cleeremans, 2003; Heun, Burkart, Wolf, & Benkert, 1998; Staresina & Davachi, 2009; Wlotko & Federmeier, 2015). Moreover, presentation rates can also alter the neural responses to those items and shift the reliance from one neural network to another (e.g., Buhusi & Meck, 2005; Foerde & Shohamy, 2011; Schultz, Tremblay, & Hollerman, 2003). Methodologically, optimal presentation settings should be determined and used in research in order to appropriately measure a given cognitive function or mental representation, including both its behavioral and neural aspects. Importantly, however, it is still unclear whether these behavioral or neural assessments, using a particular presentation rate, reflect primarily the momentary performance in the task or the underlying competence itself. This issue can affect a wide range of cognitive tasks using different presentation rate settings (e.g., response-to-stimuli intervals (RSIs), inter-trial-intervals (ITIs), or stimulus-onset asynchrony (SOA)) (Emberson, Conway, & Christiansen, 2011; Foster & Giovanello, 2017; Mechelli, Friston, & Price, 2000; Shin & Ivry, 2002). Here we aimed to systematically test, in a learning task, to what extent presentation rates affect the expression of the acquired knowledge (performance) vs. the knowledge itself (competence). To this end, we chose a procedural learning task that involves sequentially presented perceptual stimuli, which participants are required to process, respond to, and learn predictable inter-stimulus relationships – serving as an ideal model task to test the effect of presentation rate on these processes.

Procedural learning is a fundamental cognitive function that enables us to extract sequential or frequency-based regularities embedded in the environment, and contributes to the acquisition of automatic behaviors, such as skills and habits (Armstrong, Frost, & Christiansen, 2017; Fiser & Aslin, 2002; Romano, Howard, & Howard, 2010; Ullman, Earle, Walenski, & Janacsek, in press). Learning typically occurs implicitly (i.e., without conscious access either to what was learned or to the fact that learning occurred) (Cleeremans, Destrebecqz, & Boyer, 1998; Reber, 1993), although explicit (conscious) knowledge about the regularities can gradually emerge in certain cases (Cleeremans & Jiménez, 2002; Destrebecqz & Cleeremans, 2003; Jacoby, 1991; Reingold & Merikle, 1988). For example, presentation rate is a critical factor that can affect learning and the amount of explicit knowledge gained about the regularities. Previous research suggests that faster presentation rates may be more beneficial for learning than the slower ones, although results remain controversial (Arciuli & Simpson, 2011; Bogaerts, Siegelman, & Frost, 2016; Destrebecqz & Cleeremans, 2003; Emberson et al., 2011; Frensch & Miner, 1994; Soetens, Melis, & Notebaert, 2004; Willingham, Greenberg, & Thomas, 1997). Moreover, it remains unclear whether presentation rates primarily affect the expression of the acquired knowledge (performance) or the acquisition of the regularities (competence) itself. Here we systematically test to what extent slower presentation rates affect performance versus competence, including whether slower presentation rates lead to more explicit knowledge.

The effect of stimulus presentation rate on procedural learning can be experimentally tested by manipulating the Response-to-Stimulus Interval (RSI), that is, the time interval between a response and the next appearing stimulus (Willingham et al., 1997), which sets the tempo of self-paced learning. Previous studies using sequence learning tasks, such as the Serial Reaction Time (SRT) task, have shown that longer or incongruent stimulus presentation times affect learning: the longer or more incongruent the RSI, the worse the learning performance, expressed primarily in reaction times (RTs) (Destrebecqz & Cleeremans, 2003; Dominey, 1998; Frensch & Miner, 1994; Howard, Howard, Dennis, & Yankovich, 2007; Nissen & Bullemer, 1987; Soetens et al., 2004; Stadler, 1995). A possible explanation for this observation could be that shorter RSIs enable simultaneous maintenance of several consecutive items in short-term memory, promoting better learning of the sequential structure (Destrebecqz & Cleeremans, 2003; Frensch & Miner, 1994). Nevertheless, other studies showed an opposite pattern with longer inter-stimulus intervals leading to better learning, at least in some performance measures, such as accuracy index of familiarity (Arciuli & Simpson, 2011; Emberson et al., 2011). This finding may be explained by assuming that the longer RSI can promote a deeper elaboration of the learning material (e.g., the stimulus sequence), potentially leading to more explicit knowledge about the sequential structure (Destrebecqz & Cleeremans, 2003). Thus, it is possible that the stimulus presentation rate primarily affects the mode of learning (whether it promotes the formation of explicit knowledge about regularities), although this cannot be unequivocally concluded as only a few of these studies tested the amount of explicit knowledge.

Willingham et al. (1997) suggested that transfer conditions should be used to test whether participants trained with long or incongruent RSIs would show better learning performance, at least in terms of RTs, when tested with a short or consistent RSI. Using such transfer conditions, Willingham et al. (1997) showed that the longer RTs observed in the long or incongruent RSI groups primarily reflected a difficulty to express the acquired sequence knowledge but not the learning per se. Additionally, the direction of the transfer tests also proved relevant. Switching from short to long RSI led to a decrease in performance (reflected in slower RTs) compared to switching from long to short RSI (Destrebecqz & Cleeremans, 2003; Willingham et al., 1997). Importantly, however, the acquired sequence knowledge was tested only at the end of the learning phase due to the characteristics of the deterministic SRT task (RT comparison of sequence vs. random blocks is typically administered at the end of the learning phase), thus the effect of RSI on the time course of learning remains unclear. Moreover, the amount of explicit knowledge, which could have differentially affected the learning vs. expression of knowledge, was not tested in this study.

Probabilistic sequence learning tasks have multiple advantages compared to the tasks with deterministic sequences that were typically used in previous RSI studies. The Alternating Serial Reaction Time (ASRT) task is one of the most frequently used probabilistic sequence learning tasks (Howard & Howard, 1997; Kóbor, Janacsek, Takács, & Nemeth, 2017; Nemeth et al., 2010). In contrast to deterministic SRT (Nissen & Bullemer, 1987), in the ASRT, repeating sequence elements alternate with random elements throughout the task, enabling to track the time course of learning. Moreover, it has been shown that performance in the ASRT task is less affected by fatigue compared to deterministic sequence learning tasks (Pan & Rickard, 2015; Török, Janacsek, Nagy, Orbán, & Nemeth, 2017), it can therefore provide a clearer (less confounded) measure of learning.

The aim of our study was to test the effect of short vs. long RSI on the acquisition and expression of knowledge in probabilistic sequence learning. Two versions of the ASRT task were compared in healthy young adults: the fast RSI group performed the task with 120 ms RSI and the slow RSI group performed the task with 850 ms RSI in the Learning phase. The RSI lengths were determined based on previous studies (Howard & Howard, 1997; Kóbor et al., 2017; Nemeth et al., 2010; Willingham et al., 1997). To test whether fast vs. slow RSI affected mainly performance or competence, the two groups performed both RSI versions of the task 24 hours later in the Testing phase. This retention period was chosen to ensure that well-consolidated knowledge is tested, thus the effect of RSI change is not confounded with further learning effects in the Testing phase (Kóbor et al., 2017; Nemeth & Janacsek, 2011). Overall, our study addresses three questions: (1) Does fast vs. slow RSI affect performance in the Learning phase? (2) Does fast vs. slow RSI affect mainly the expression of knowledge (performance) or learning per se (competence)? (3) How does the stimulus presentation rate affect the amount of explicit knowledge gained in the task?

First, we hypothesized that the RSI length affects the performance in the Learning phase: namely, the slow RSI group will show weaker learning scores compared to the fast RSI group (Frensch & Miner, 1994; Soetens et al., 2004). Second, we expected that the RSI length affects mainly the expression of but not the acquired knowledge itself: namely, the groups tested with similar RSI will exhibit similar learning scores in the Testing phase, irrespective of the RSI during learning (Howard et al., 2007; Willingham et al., 1997). Third, we hypothesized that the slow RSI group will exhibit greater explicit sequence knowledge compared to the fast RSI group (Destrebecqz & Cleeremans, 2003).

## 2 Materials and methods

### 2.1 Participants

Seventy-nine persons (68 females/11 males) took part in the experiment. They were between the ages of 19 to 30 (*M*_Age_ = 22.00 years, *SD*_Age_ = 1.88 years). They were university students from Budapest, Hungary (*M*_Years of education_ = 14.67 years, *SD*_education_ = 1.6 years). Handedness was measured by the Edinburgh Handedness Inventory (Oldfield, 1971), the Laterality Quotient of the sample varied between −80 and 100 (−100 means complete left-handedness, 100 means complete right-handedness, *M*_LQ_ = 47.30, *SD*_LQ_ = 47.54). Participants were randomly assigned to one of four experimental groups (see Procedure and Figure 1.B). None of the participants reported history of developmental, psychiatric, neurological or sleep disorders and they had normal or corrected-to-normal vision. They performed in the normal range on standard neuropsychological tests of short-term and working memory, and their cognitive performance did not differ (Digit span task: *M* = 6.06, *SD* = 1.03, Counting span task: *M* = 3.76, *SD* = 1.16) (Janacsek & Nemeth, 2013). Before the assessment, all participants gave signed informed consent and received course credit for participation. The study was approved by the Institutional Review Board of Eötvös Loránd University, Hungary.

**Figure 1.**
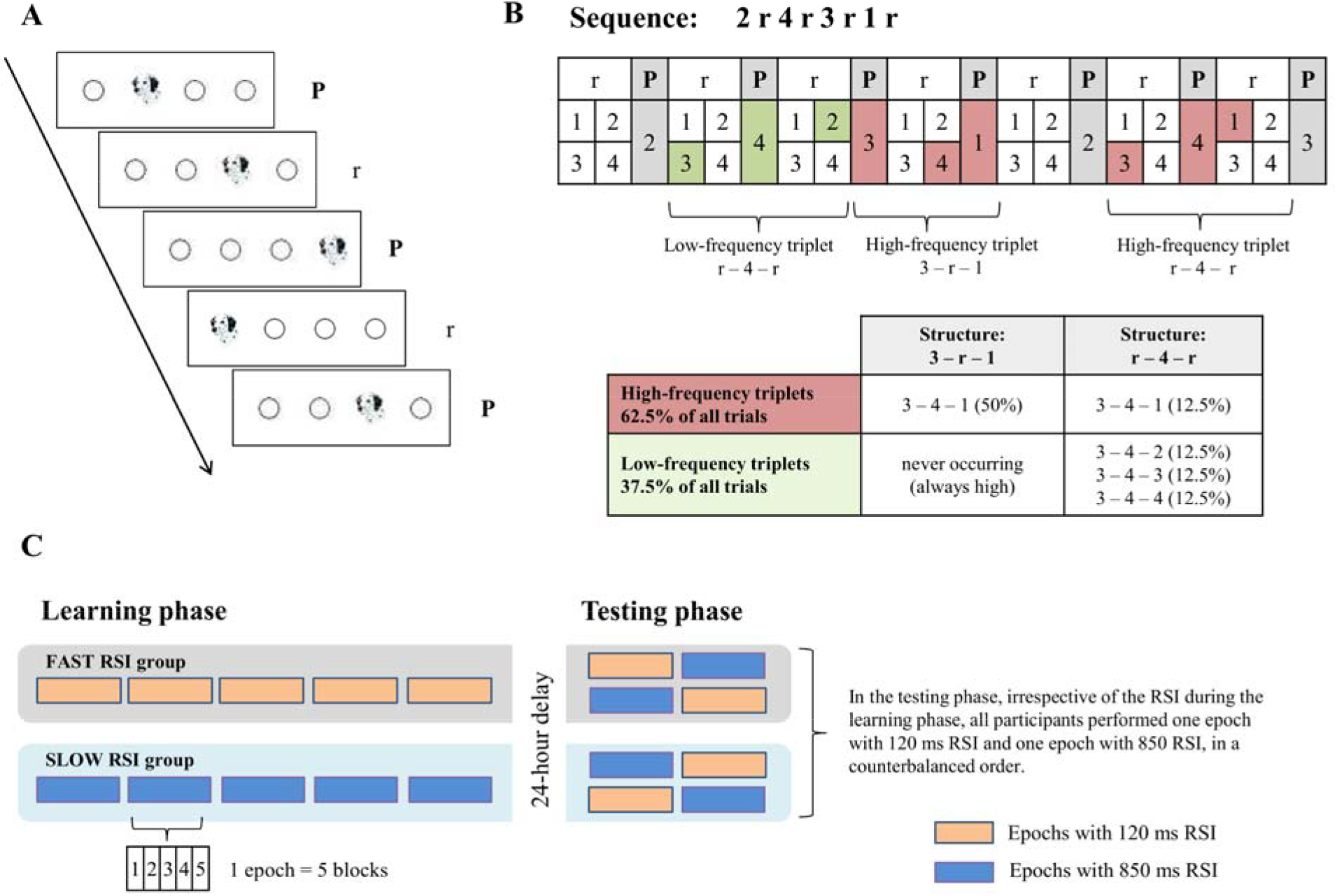
Design of the experiment. (A) In the Alternating Serial Reaction Time (ASRT) task participants were asked to respond to the stimuli (dog’s head) appearing on one of four locations, and press the corresponding key as fast and as accurately as they could. Unbeknownst to them, every second stimulus was part of a repeated alternating sequence (P – pattern) and every second stimulus was random (r – random). (B) As a result of the alternation of pattern and random trials, there were more probable and less probable combinations of three consecutive stimuli (high-frequency and low-frequency triplets, respectively). Learning was measured as differences in responses to high- vs. low-frequency triplets, both in reaction time and accuracy. (C) The experiment consisted of two sessions, separated by a 24-hour delay. On Day 1 (*Learning phase*) participants were randomly assigned to one of two groups: the fast RSI group performed 25 blocks (5 epochs) of the ASRT task with 120 ms response-to-stimulus interval (RSI), the slow RSI group performed 25 blocks (5 epochs) of the task with 850 ms RSI. On Day 2 (*Testing phase*), all participants performed 10 blocks (2 epochs) of the task: 5 blocks with the fast (120 ms) and 5 blocks with the slow (850 ms) version of the task, in counterbalanced order.

### 2.2 Tasks

#### 2.2.1 Alternating Serial Reaction Time (ASRT) Task

Learning was measured by the ASRT task (Howard & Howard, 1997; Nemeth et al., 2010). In this task, a stimulus (a dog’s head) appeared in one of four horizontally arranged empty circles on the screen and participants had to press the corresponding button as quickly and accurately as they could when the stimulus occurred. The computer was equipped with a keyboard with four heightened keys (Z, C, B, M on a QWERTY keyboard), each corresponding to the circles in a horizontal arrangement. The stimulus remained on the screen until the participant pressed the correct button. The next stimulus appeared after a 120 or 850 ms response-to-stimulus-interval (RSI) (for more details on the presentation rate see Procedure). The task was presented in blocks of 85 stimuli: unbeknownst to the participants, after the first five warm-up trials consisting of random stimuli, an 8-element alternating sequence was presented ten times (e.g., 2r4r3r1r, where each number represents one of the four circles on the screen and r represents a randomly selected circle out of the four possible ones). Due to the alternating sequence structure, some triplets (i.e., runs of three consecutive events) occurred more frequently than others did. Following previous studies, we refer to the former as *high-frequency triplets* and the latter as *low-frequency triplets*. Note that due to the higher occurrence probability, the final event of high-frequency triplets was more predictable from the initial event of the triplet compared to the low-frequency triplets. For each stimulus/event, we determined whether it was the last element of a high- or low-frequency triplet.

**Figure 2.**
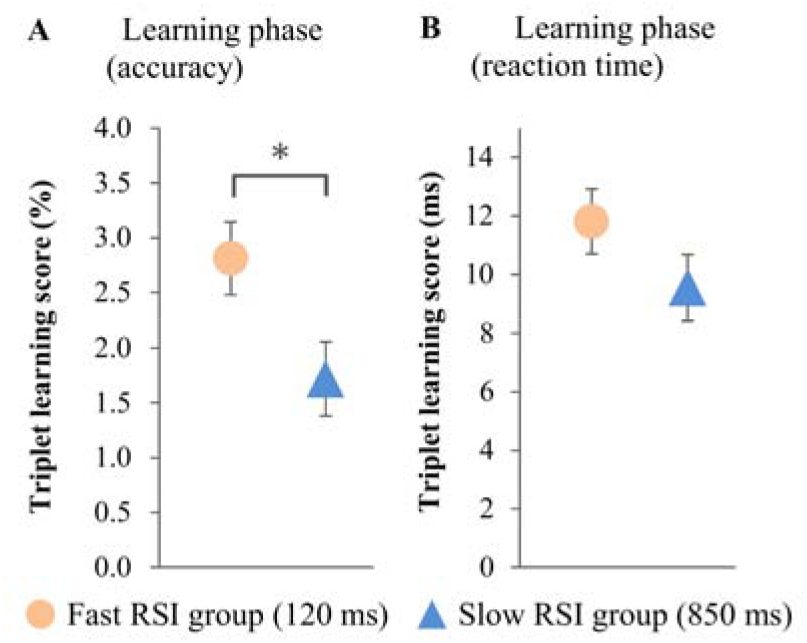
Triplet learning scores in the Learning phase, measured by accuracy (A) and reaction time (B). The fast RSI group showed significantly larger triplet learning scores compared to the slow RSI group, at least in terms of accuracy. The error bars represent the standard error of means (SEM). ** *p* < 0.01; * *p* < 0.05; + *p* < 0.1.

#### 2.2.2 Inclusion-Exclusion Task

We administered the Process Dissociation Procedure (PDP; Jacoby, 1991) using an Inclusion-Exclusion Task to test whether the participants gained explicit or conscious knowledge about the statistical regularities underlying the ASRT task. Before performing this task, participants were informed that the order of the stimulus appearance followed a hidden sequence. First, we asked them to generate this sequence (*Inclusion condition*) using the same response buttons as the ones used during the ASRT task. They were told to rely on their intuitions if they were unsure about the sequence. They performed four runs of this inclusion condition. Then they were asked to generate a sequence of responses that was different from the learned ASRT sequence (*Exclusion condition*). Due to the alternating ASRT sequence structure, some combinations of the stimuli could never occur in the task (such as systematic sequences across pattern *and* random trials, e.g., 12341234; systematic alterations, e.g., 12123434; systematically rare or frequent occurrences of the same stimulus, e.g., 11111233334; or a longer stimulus stream without a specific stimulus, e.g., 123321123). Since participants are asked to generate a *realistic* ASRT sequence that they learned (in the Inclusion condition) or that is different from the learned one (in the Exclusion condition), they are asked to avoid these systematic combinations that could never occur in *any* ASRT sequence (not just the learned one). They performed four runs of this condition as well. In both conditions each run was finished after 24 key presses were made (Kóbor et al., 2017).

**Figure 3.**
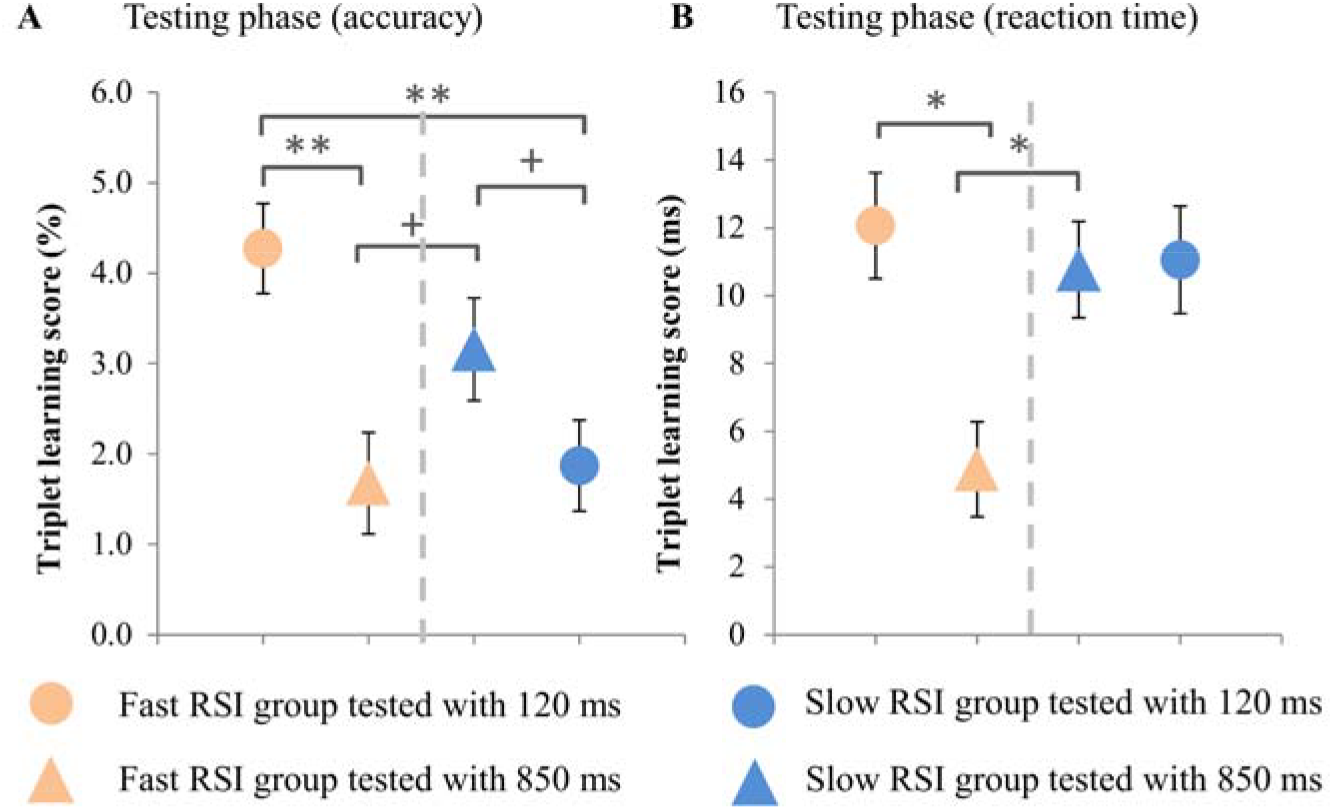
Triplet learning scores in the Testing phase, measured by accuracy (A) and reaction time (B). The colors represent the groups that *learnt* with different presentation rates (orange – fast RSI group, blue – slow RSI group), and the shapes represent different presentation rates during *testing* (circle – tested with 120 ms, triangle – tested with 850 ms). The triplet learning scores measured by accuracy shows that participants exhibited smaller learning scores when tested with a different presentation rate than the one encountered during learning compared to when they were tested with the same presentation rate. In terms of the triplet learning scores measured by reaction time, the group that learnt with the faster presentation rate showed weaker learning scores when tested with the slower presentation rate. The error bars represent SEM. ** *p* < 0.01; * *p* < 0.05; + *p* < 0.1.

According to the PDP, in the inclusion condition successful performance can achieved by both the implicit and explicit knowledge, but in the exclusion condition the implicit and explicit knowledge compete with each other, only the explicit, conscious knowledge can lead to generate a different sequence of key presses. To analyze it, we calculated the percentage of producing high-frequency triplets in the inclusion and exclusion conditions separately. Then we compared these scores to chance level. In this case it is 25%, because out of the 64 potential triplets that participants can generate, only 16 are high-frequency ones. If participants generated more high-frequency triplets in the inclusion condition than it would have been expected by chance, it can indicate either implicit or explicit knowledge about the statistical regularities. But if they generated more high-frequency triplets in the exclusion condition than it would have been expected by chance, it can indicate implicit knowledge and lack of conscious control over their explicit knowledge (Kóbor et al., 2017).

### 2.3 Procedure

There were two sessions in the experiment: a Learning phase and a Testing phase separated by a 24-hour long delay. On the first day (*Learning phase*) we administered the ASRT task. Participants were not given any information about the regularity that was embedded in the task (Nemeth et al., 2010). They were informed that the main aim of the study was to test how extended practice affected performance on a simple reaction time task. Therefore, we emphasized performing the task as accurately and as fast as they could. Between blocks, the participants received feedback about their overall accuracy and reaction time presented on the screen, and then they had a rest period of between 10 and 20 s before starting the next block.

Participants were randomly assigned to one of two groups on Day 1: 40 participants performed the fast RSI version, and 39 performed the slow RSI version of the ASRT task. In the fast version of the task, the RSI was 120 ms, while in the slow version, the RSI was 850 ms. The ASRT consisted of 25 blocks on Day 1. In the fast version one block took about 1-1.5 min, and in the slow version one block took about 1.5-2.5 min, therefore the fast version of the task took approximately 30 min, and the slow one approximately 45 min.

On Day 2, all participants performed 10 blocks of the ASRT task: 5 blocks with the fast and 5 blocks with the slow version of the task, in counterbalanced order (See Figure 1.B). For potential order effects, see Supplementary results. The fast blocks took about 5-8 mins, and the slow ones approximately 10-13 min, the Testing phase totally was 15-22 min long.

For each participant, one of the six unique permutations of the four possible ASRT sequence stimuli was selected in a pseudorandom manner, so that the six different sequences based on a permutation rule were used across participants (Howard & Howard, 1997; Nemeth et al., 2010). On Day 2, participants were tested with the same ASRT sequence as the one they learnt on Day 1. Note that the ASRT task has good test-retest reliability, and it is therefore a suitable tool to assess learning, especially in multiple sessions (Stark-Inbar, Raza, Taylor, & Ivry, 2016; West, Vadillo, Shanks, & Hulme, 2017).

After performing the ASRT task on Day 2, we tested the amount of explicit knowledge the participants acquired about the task with a short questionnaire and the Inclusion-Exclusion Task. The short questionnaire (Nemeth et al., 2010; Song, Howard, & Howard, 2007) included two questions: „Have you noticed anything special regarding the task?” and „Have you noticed some regularity in the sequence of stimuli?”. None of the participants reported noticing the hidden regularity in the task. The results of the Inclusion-Exclusion Task are discussed in the Results section.

### 2.4 Statistical analysis

We followed the standard data processing and analysis protocols of previous ASRT studies (Kóbor et al., 2017; Nemeth, Janacsek, Király, et al., 2013). Based on these protocols, epochs of five blocks were analyzed instead of single blocks (thus, Epoch 1 corresponds to Blocks 1-5, Epoch 2 corresponds to Blocks 6-10, etc.). We calculated mean accuracy for all trials and median RTs for correct responses only for each participant and each epoch, separately for high- and low-frequency triplets. Accumulating behavioral evidence indicates that participants respond increasingly accurately and faster to high-frequency triplets compared with low-frequency ones as the ASRT task progresses, revealing procedural learning (e.g. Howard & Howard, 1997; Nemeth, Janacsek, & Fiser, 2013). Therefore, triplet learning scores were calculated as a difference in RT/accuracy for high- vs. low-frequency triplets (for RTs: low- minus high-frequency; for accuracy: high- minus low-frequency). Higher learning scores indicate better learning/performance in the task.

Triplet learning scores were submitted to a series of mixed-design analyses of variance (ANOVAs) on Day 1 and Day 2 (for details see the Results section). In all ANOVAs, the Greenhouse-Geisser epsilon (ε) correction (Greenhouse & Geisser, 1959) was used when necessary. Original *df* values and corrected *p* values are reported (if applicable) together with partial eta-squared *η*_*p*_^2^ as the measure of effect size. LSD (Least Significant Difference) tests for pair-wise comparisons were used to control for Type I error.

Finally, we analyzed the Inclusion-Exclusion Task following the procedures described in previous ASRT studies (Horvath, Torok, Pesthy, Nemeth, & Janacsek, 2018; Kóbor et al., 2017). To ensure that all participants followed the instructions in the two conditions, we examined and excluded those runs in which participants generated systematic combinations of stimuli explained in the task description, and thus, they did not follow the instructions. From all 632 runs we excluded 68, so the analysis contains the 89.24% of the answers. As a condition, the amount of the high-frequency triplets was counted. This measure was also submitted to a mixed design ANOVA to test if participants gained explicit triplet knowledge.

## 3 Results

### 3.1 Does the length of the RSI affect performance in the Learning phase?

To test whether the length of the RSI affects performance in the Learning phase, we compared the performance of the fast (120 ms) and slow (850 ms) RSI groups on Day 1. The triplet learning scores, separately for accuracy and reaction time, were analyzed by a mixed design ANOVA with EPOCH (1-5) as a within-subject factor, and LEARNING RSI (fast vs. slow RSI) as a between-subject factor.

#### Accuracy

The ANOVA revealed significant triplet learning [indicated by the significant INTERCEPT: *F*_(1, 77)_ = 90.113, *p* < 0.001, *η*_*p*_^2^ = 0.539] such that participants were more accurate on high-than on low-frequency triplets. As the task progressed, the triplet learning scores increased [indicated by the significant main effect of EPOCH: *F*_(4, 74)_ = 5.265, *p* < 0.001, *η*_*p*_^2^ = 0.064]. The two groups showed different triplet learning scores overall [significant main effect of LEARNING RSI: *F*_(1, 77)_ = 5.660, *p* = 0.020, *η*_*p*_^2^ = 0.068]: the fast RSI group greater triplet learning scores than the slow RSI group [2.8% vs. 1.7%, respectively], suggesting that the length of the RSI affected triplet learning scores on Day 1. The time course of learning did not differ between the two groups [EPOCH x LEARNING RSI: *F*_(4, 74)_ = 0.271, *p* = 0.897, η_p_ ^2^ = 0.004*η*_*p*_^2^ = 0.004].

#### Reaction Time

A similar ANOVA was conducted for reaction time data. The ANOVA revealed significant triplet learning scores [showed by the significant INTERCEPT: *F*_(1, 77)_ = 181.957, *p* < 0.001, *η*_*p*_^2^ = 0.703] such that RTs were faster on high-than on low-frequency triplets. As the task progressed, the participants’ triplet learning scores increased [indicated by the significant main effect of EPOCH: *F*_(4, 74)_ = 11.768, *p* < 0.001, *η*_*p*_^2^ = 0.133]. The two groups did not differ in overall triplet learning [main effect of LEARNING RSI: *F*_(1, 77)_ = 2.024, *p* = 0.159, *η*_*p*_^2^ = 0.026], however, the time course of learning was marginally different between the groups [EPOCH x LEARNING RSI interaction: *F*_(4, 74)_ = 2.113, *p* = 0.079, *η*_*p*_^2^ = 0.027]. The LSD post-hoc test revealed that the two groups’ triplet learning scores were similar in the first 3 epochs [all *p*s > 0.344], but the fast RSI group showed greater triplet learning scores than the slow RSI group in Epoch 4 and 5 [all *p*s < 0.037]. These results suggest that the length of RSI had an impact on triplet learning scores on Day 1.

### 3.2 What does the RSI length affect: the acquired knowledge (competence) or just the expression of knowledge (performance)? Results of the Testing phase

To test whether the RSI length affects the acquired knowledge or the expression of that knowledge, we analyzed the ACC and RT data of the Testing phase. Independent of the RSI during the Learning phase, here all participants were tested with both RSI length (120 and 850 ms), in a counterbalanced order. We conducted a mixed design ANOVA for TEST RSI (tested with 120 vs. 850 ms RSI, independently of whether it was the first or the second epoch) as a within-subject factor, and LEARNING RSI (fast vs. slow RSI) as a between-subject factor.

#### Accuracy

The ANOVA revealed no significant main effect of TEST RSI [*F*_(1, 77)_ = 1.774, *p* = 0.187, *η*_*p*_^2^ = 0.023], or LEARNING RSI [*F* = 0.670, *p* = 0.416, *η*_*p*_^2^ =0.009]. The TEST RSI x LEARNING RSI interaction was, however, significant [*F*_(1, 77)_ = 14.086, *p* < 0.001, *η*_*p*_^2^ = 0.155], suggesting that the performance in the Testing phase depended on both the RSI length during learning and during testing. Post hoc tests revealed that, when *tested* with 120 ms RSI, the group that *learnt* with 120 ms showed significantly greater learning scores than the group that *learnt* with 850 ms [4.3% vs. 1.9%, respectively, *p* = 0.001]. An opposite pattern was observed when *tested* with 850 ms RSI: the group that *learnt* with 120 ms demonstrated marginally smaller learning scores than the group that *learnt* with 850 ms [1.7 % vs. 3.2 %, respectively, *p* = 0.062]. The group that *learnt* with 120 ms RSI showed greater learning scores when *tested* with 120 ms RSI compared to the 850 ms RSI [*p* = 0.001]. An opposite trend was observed for the group that *learnt* with 850 ms RSI: they showed smaller learning scores when *tested* with 120 vs. 850 ms RSI [*p* = 0.093]. These results suggest the presentation rates primarily affected the expression of the acquired knowledge (performance) rather than competence as when tested with a different presentation rate than the one encountered during learning, participants exhibited smaller learning scores compared to when they were tested with the same presentation rate. Neither presentation rate (fast or slow RSI) alone led to a superior performance compared to the other; instead, a congruency effect was present as both groups exhibited better learning scores when the same RSI length was used during the Learning and the Testing phases. The remaining main effects and interactions did not reach significance [all *p*s ≥ 0.187].

#### Reaction Time

For the RT learning scores, the ANOVA revealed that, independently from the RSI of the Learning phase, all groups showed higher learning scores when *tested* with 120 ms RSI compared to 850 ms RSI [indicated by the significant main effect of TEST RSI: *F*_(1, 77)_ = 6.568, *p* = 0.012, *η*_*p*_^2^ = 0.079]. The ANOVA also revealed a significant TEST RSI x LEARNING RSI interaction [*F*_(1, 77)_ = 5.613, *p* = 0.020, *η*_*p*_^2^ = 0.068]. The post hoc tests revealed that, when *tested* with 120 ms, both groups that *learnt* with 120 vs. 850 ms RSI showed similar learning scores [12.069 ms vs. 11.058 ms, respectively, *p* = 0.650]. In contrast, when *tested* with 850 ms, the group that *learnt* with 120 ms RSI showed significantly smaller learning scores than the group that *learnt* with 850 ms [4.881 ms vs 10.776 ms, respectively, *p* = 0.004]. The group that *learnt* with 120 ms RSI exhibited greater learning scores when *tested* with 120 ms RSI compared to 850 ms RSI [12.069 ms vs. 4.881 ms, respectively, *p* = 0.001], whilst the group that *learnt* with 850 ms RSI, showed similar learning scores when tested with 120 vs. 850 ms RSI [11.058 ms vs. 10.776 ms, respectively, *p* = 0.892]. These results suggest that the slower presentation rate (850 ms RSI) during learning primarily affected the expression of the acquired knowledge (performance) rather than competence as 1) when tested with the faster presentation rate (120 ms RSI), the participants’ learning scores were comparable to that of the group that encountered the faster presentation rate during learning as well. 2) Additionally, the group that learnt with the faster presentation rate showed weaker learning scores when tested with the slower presentation rate.

### 3.3 Does RSI affect the explicit knowledge gained in the task?

Overall, the generated percentage of high-frequency triplets differed significantly from chance level both in the inclusion and exclusion conditions [one-sample t-tests, inclusion condition: *t*_(76)_ = 9.441, *p* < 0.001; exclusion condition: *t*_(74)_ = 8.093, *p* < 0.001]. To test whether RSI affects how much explicit knowledge participants gain in the task, a mixed design ANOVA was conducted with CONDITION (inclusion vs. exclusion) as a within-subject factor, and LEARNING RSI (fast vs. slow) as a between-subject factor. The main effect of CONDITION did not reach significance [*F*_(1, 69)_ = 0.696, *p* = 0.407, *η*_*p*_^2^ = 0.010]: participants generated a greater percentage of high-frequency triplets than it would have been expected by chance both in the inclusion and exclusion conditions [7.5% and 6.6% above chance level, respectively]. These results indicate that participants could not exclude their acquired knowledge when they were instructed to do so. The remaining main effects and interactions did not reach significance [*p* ≥ 0.735], suggesting no group differences in the amount of acquired knowledge. In sum, based on this analysis, the triplet knowledge in the ASRT task remained implicit in all groups.

## 4 Discussion

In our study, we systemically tested the effect of presentation rate on the acquisition and expression of knowledge in probabilistic sequence learning. Consistent with our hypothesis, we found that the slower presentation rate led to weaker learning performance in the Learning phase. Importantly, however, the results of the Testing phase showed that this effect was primarily due to the expression of the acquired knowledge and not the learning itself. Thus, the slower presentation rate affected performance rather than competence, in terms of both reaction times and accuracy. Finally, the acquired knowledge remained implicit in both groups, regardless of the presentation rate during learning.

Previous studies suggested that the slower presentation rate lead to weaker learning performance, at least in reaction times, because it impedes the simultaneous maintenance of several consecutive items of the sequence, hindering the learning of the sequential structure (Destrebecqz & Cleeremans, 2003; Dominey, 1998; Frensch & Miner, 1994; Soetens et al., 2004; Stadler, 1995). In other words, at a slower presentation rate, the consecutive items are further apart from one another in time, making it less likely to be represented and bind together in a local short-term storage or cache (Janacsek & Nemeth, 2013, 2015). This explanation, however, does not seem likely as we found that when learnt with the slow RSI but tested with the fast RSI, participants’ performance was comparable to that of the group that encountered the fast RSI during learning as well. Additionally, the group that learnt with the fast RSI showed weaker performance when tested with the slow RSI. Importantly, accuracy measures also suggest that presentation rates primarily affect the expression of knowledge. When tested with a different RSI than the one encountered during learning, participants exhibited weaker learning performance compared to when they were tested with the same RSI. Neither presentation rate alone led to a superior performance compared to the other; instead, a congruency effect was present as both groups exhibited better learning performance when the same RSI length was used during the Learning and the Testing phases (see also Supplementary results I). Thus, it appears that presentation rates primarily affect the expression of knowledge (performance) and not the sequence learning itself (competence). For reaction time measures, slowing down the presentation rates seem to have a more adverse effect compared to speeding up the presentation rates, while for accuracy measures, expression of knowledge is better when the presentation rate is congruent with the original one encountered during learning, irrespective of the presentation rate (fast or slow) itself.

According to an alternative interpretation, the slower presentation rates may increase the chance of gaining more explicit knowledge about the sequence structure as there is more time for a deeper elaboration of the learning material (Destrebecqz & Cleeremans, 2003). However, this does not seem to be the case in our study as we showed that knowledge remained implicit in both groups, regardless of the presentation rate during learning. It is important to note that the sequence structure may affect whether and how much explicit knowledge participants gain during learning. Previous studies showed that deterministic sequences are more likely to become explicit during learning compared to probabilistic sequences (Kóbor et al., 2017; Nemeth et al., 2010). Thus, it is possible that presentation rates affect the mode of learning (whether it promotes the formation of explicit knowledge about regularities) differently for deterministic vs. probabilistic sequences.

Why did stimulus presentation rates affect the expression of knowledge? One possibility is that faster presentation rates lead to response facilitation in that consecutive stimuli are closer to one another and thus the representation of the previous stimuli may be still activated in the time window when response is made to the current stimulus. Since the previous stimuli predict the current stimulus with a certain probability, their activation in this time window may facilitate the response to the current stimulus, which may be reflected in faster RTs and/or higher accuracy (Janacsek, Borbély-Ipkovich, Nemeth, & Gonda, 2018; Takács et al., 2018). This response facilitation may be greater for more predictable stimulus combinations (such as the high-frequency triplets in our study) compared to the less predictable ones. As more time elapses between subsequent stimuli, response facilitation may be weaker for slower presentation rates, resulting in smaller differences in responding to more and less predictable stimuli (which may be more pronounced as more time elapses in a given block, i.e., at the later part of each block, see Methods and Supplementary results II). Importantly, however, based on our results of the Testing phase, this weaker response facilitation affected only the expression of knowledge and not the sequence learning (e.g., the binding of consecutive sequence stimuli) itself. Thus, it appears that learning predictable regularities of sequential patterns can occur even with slower presentation rates (Arciuli & Simpson, 2011; Bogaerts et al., 2016; Destrebecqz & Cleeremans, 2003; Emberson et al., 2011), even if it is not simultaneously reflected in performance measures to the same amount as in faster presentation rates. Further research seems warranted to determine the upper limit of the presentation rate that still enables the acquisition of temporally distributed sequential regularities, which likely will depend on various factors such as sequence predictability (e.g., deterministic vs. probabilistic sequences) and sequence length.

In summary, our study highlights that the expression of knowledge can dissociate from the learning itself: in procedural learning, slower stimulus presentation rates can lead to weaker performance, masking the acquired knowledge. This suggests that an optimal performance could be achieved if the environment or task enables a faster pace. Our findings can have important implications both for research and practice (e.g., in education, language learning, sports as well as in clinical fields). From a practical perspective, for instance, when learning a foreign language or mastering sports, students might show better performance if speeded processing and responding is required. In clinical neuroscience, some patient populations may show generally slower responses in self-paced tasks compared to healthy participants, which can mask their acquired knowledge, possibly leading to incorrect interpretations. Finally, from a research perspective, our findings highlight that more well-designed tasks and test protocols are needed that are able to disentangle competence from performance, in multiple domains of cognition such as learning, memory, decision-making, perception and language.

## Supporting information

Supplemental results

## Acknowledgments

We are thankful to Zsuzsa Londe for her useful comments on the previous draft of the manuscript. This research was supported by the National Brain Research Program (project 2017-1.2.1-NKP-2017-00002); Hungarian Scientific Research Fund (OTKA PD 124148, to KJ; NKFIH-OTKA K 128016, to DN); Janos Bolyai Research Fellowship of the Hungarian Academy of Sciences (to KJ). DN appreciates the support from IMÉRA.

