## Supplemental results for "Stimulus presentation rates affect performance but not the acquired knowledge – Evidence from procedural learning"

### Supplementary results I: Does the order of testing influence performance in the Testing phase?

As presented in the main text, on Day 2, all participants performed 10 blocks of the ASRT task: 5 blocks with the fast and 5 blocks with the slow version of the task, in counterbalanced order. Consequently, half of the participants were first tested with the same timing as the one during the Learning phase (termed 'congruent-first'), while the other half of the participants were first tested with the other timing than the one during the Learning phase (termed 'incongruent-first'). Based on previous research, it is possible that participants exhibit weaker performance when they are tested with the incongruent timing first (Dominey, 1998; Stadler, 1995). Additionally, combined with the main hypothesis presented in the manuscript, it may be expected that this order effect interacts with the length of RSI. Namely, the incongruent-first testing may impair performance especially when it means increasing RSI from the Learning to the Testing phase (i.e., the 120 ms RSI group performs worse when tested with 850 ms RSI first, compared to 850 ms RSI group when tested with the 120 ms RSI first).

We conducted mixed design ANOVAs with EPOCH (1 vs. 2) as a within-subject factor, and LEARNING RSI (fast vs. slow RSI) and TESTING ORDER (2: congruent-first vs. incongruent-first) as between-subjects factors.

**Accuracy** – The ANOVA revealed a marginally significant main effect of EPOCH [$F_{(1, 75)} = 3.693$, $p = 0.058$, $\eta_p^2 = 0.047$] and a significant EPOCH x TESTING ORDER interaction: $F_{(1, 75)} = 14.214$, $p < 0.001$, $\eta_p^2 = 0.159$]. The *LSD post-hoc* tests revealed that participants showed better learning scores when they were tested with the same RSI as during learning (similar to the accuracy results for Hypothesis 2 in the main text). Namely, in Epoch 1, the participants who were tested with the congruent RSI first performed significantly better than the ones tested with the incongruent RSI first [3.5 % vs. 1.0 %, respectively, $p = 0.001$]. In Epoch 2, those participants who were tested with the incongruent RSI first tended to show better learning scores than the ones who were tested with the congruent RSI first [2.5 % vs. 3.9 %, respectively, $p = 0.078$]. Note that for those participants who were tested with the incongruent RSI first, the RSI in Epoch 2 was the same as during learning, thus showing a congruency effect.

Additionally, the LEARNING RSI x TESTING ORDER interaction was also significant [$F_{(1, 75)} = 5.911$, $p = 0.017$, $\eta_p^2 = 0.073$]. The LSD *post hoc* test revealed that, when tested with the congruent RSI first, the group that learnt with 120 ms RSI (i.e., fast RSI group) showed greater learning scores than the group that learnt with 850 ms RSI (i.e., slow RSI group) [3.9 % vs. 2.1 %, respectively, $p = 0.023$], in accordance with the results of the Learning phase in the main text. When tested with the incongruent RSI first, the two groups' learning scores were similar [$M_{120\ \text{Incongruent-first}} = 2.1$ % vs $M_{850\ \text{Incongruent-first}} = 2.9$ %, $p = 0.268$]. Within the fast RSI group, those who were tested with the congruent RSI first showed greater learning scores than those who were tested with the incongruent RSI first [$M_{120\ \text{Congruent-first}} = 3.9$ % vs $M_{120\ \text{Incongruent-first}} = 2.1$ %, $p = 0.017$]. Within the slow RSI group, there was no difference between those who were tested with the congruent vs. incongruent RSI first [$M_{850\ \text{Congruent-first}} = 2.1$ % vs $M_{850\ \text{Incongruent-first}} = 2.9$ %, $p = 0.316$].

These results together highlight that 1) greater learning scores could be obtained when tested with the same presentation rate as during learning (congruency effect), irrespective of the presentation rate, and 2) the order of testing had a larger effect on the performance of the fast RSI group compared to the slow RSI group (see Figure S1A). The remaining main effects and interactions did not reach significance [$p \geq 0.171$].

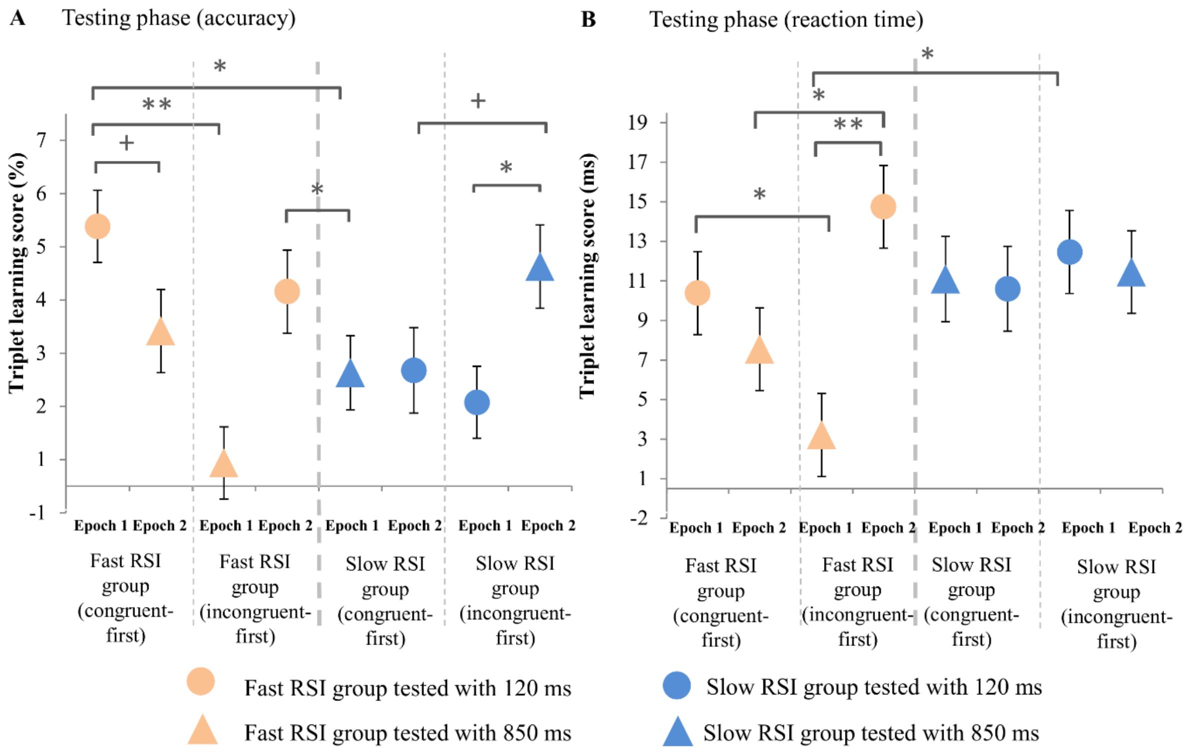


**Figure S1. The effect of testing order on performance.** The accuracy (A) and reaction time (B) triplet learning scores are presented in the Testing phase. The colors represent the groups that *learnt* with different presentation rates (orange – fast RSI group, blue – slow RSI group), and the shapes represent different presentation rates during *testing* (circle – tested with 120 ms, triangle – tested with 850 ms). Half of the participants were first tested with the same presentation rate as during learning (congruent-first), and the other half were first tested with the different presentation rate than during learning (incongruent-first). Overall, all participants were tested with both presentation rates in a counterbalanced order (shown in Epoch 1 and 2). Both in accuracy and reaction time measures the fast RSI group performed weaker when they were tested incongruent-first, compared to the slow RSI group, which was less affected by the changes in presentation rates. The error bars represent the standard error of means (SEM). ** $p < 0.01$; * $p < 0.05$; + $p < 0.1$.

**Reaction time** – A similar ANOVA conducted on the RT data revealed a significant EPOCH x TESTING ORDER interaction [$F_{(1,\ 75)} = 5.849$, $p = 0.018$, $\eta_p^2 = 0.072$], and a trend level EPOCH x LEARNING RSI interaction [$F_{(1,\ 75)} = 3.172$, $p = 0.079$, $\eta_p^2 = 0.041$]. Both interactions were qualified by the significant EPOCH x LEARNING RSI x TESTING ORDER interaction [$F_{(1,\ 75)} = 6.770$, $p = 0.011$, $\eta_p^2 = 0.083$]. The fast RSI group showed significantly smaller learning scores when tested with the incongruent (850 ms) RSI compared to the congruent (120 ms) RSI, either if it was in the first [$M_{120\ Incongruent\text{-}first\_Epoch1}$ = 2.713 ms vs. $M_{120C\_epoch1}$ = 9.888 ms, $p = 0.018$] or in the second epoch [$M_{120\ Incongruent\text{-}first\_Epoch2}$ = 14.250 ms vs. $M_{120\ Congruent\text{-}first\_Epoch2}$ = 7.050 ms, $p = 0.017$]. If we compare the congruent and incongruent conditions *within* groups, this result was more prominent in those participants who were tested with the incongruent RSI first [$M_{120\ Incongruent\text{-}first\_Epoch1}$ = 2.713 ms vs. $M_{120\ Incongruent\text{-}first\_Epoch2}$ = 14.250 ms, $p < 0.001$]. In contrast, there was no significant difference in the learning scores of the two epochs in those participants who were tested with the congruent RSI first [$M_{120\ Congruent\text{-}first\_Epoch1}$ = 9.888 ms vs. $M_{120\ Congruent\text{-}first\_Epoch2}$ = 7.050 ms, $p = 0.322$]. For the slow RSI group, there was no significant difference neither between the congruent and incongruent conditions, nor between their order of testing [$M_{850\ Congruent\text{-}first\_Epoch1}$ = 10.592 ms, $M_{850\ Congruent\text{-}first\_Epoch2}$ = 10.105 ms, $M_{850\ Incongruent\text{-}first\_Epoch1}$ = 11.963 ms, $M_{850\ Incongruent\text{-}first\_Epoch2}$ = 10.950 ms; $p$s ≥ 0.650]. The remaining main effects and interactions did not reach significance [all $p$s ≥ 0.121].

Overall, the results of RT data show that the order of testing primarily affected the performance of the fast RSI group but not that of the slow RSI group (see Figure S1B).

### Supplementary results II: Testing the within block position effect

The within block position effect refers to the phenomenon that during a longer reaction time task arranged into blocks (e.g., several seconds or minutes) participants show different performance when the first vs. second halves of the blocks are compared (Pan & Rickard, 2015; Torok, Janacsek, Nagy, Orban, & Nemeth, 2017). Such differences have been shown in some cases, such as in children or older adults, or in psychiatric/neurological conditions (Gamble et al., 2014; Nemeth, Janacsek, & Fiser, 2013; Nemeth, Janacsek, Király, et al., 2013). For example, patients with Mild Cognitive Impairments showed better triplet learning in the second halves of the blocks (Nemeth, Janacsek, Király, et al., 2013), while patients with Parkinson's disease showed the opposite pattern (Gamble et al., 2014). Here we aimed to test whether the length of RSI affects these within block position effects. Therefore, we performed the mixed design ANOVAS presented in the main text with the HALF-BLOCK as an additional within-subject factor. Note that here we do not report those effects that are already included in the main ANOVA in the Results section (e.g., main effect of LEARNING RSI or LEARNING RSI x EPOCH interaction), but only those main effects and interactions that involve the HALF-BLOCK factor.

#### *Does the length of the RSI differentially affect performance in the first vs. second halves of the blocks in the Learning phase?*

**Accuracy** – We found that participants showed greater triplet learning in the second halves of the blocks than in the first halves [shown by the significant main effect of HALF-BLOCK: $F_{(1,\ 77)} = 4.817$, $p = 0.031$, $\eta_p^2 = 0.059$; 2.6% vs. 1.9%, respectively; Figure S2A]. The remaining interactions involving HALF-BLOCK did not reach significance [all $p$s > 0.467].

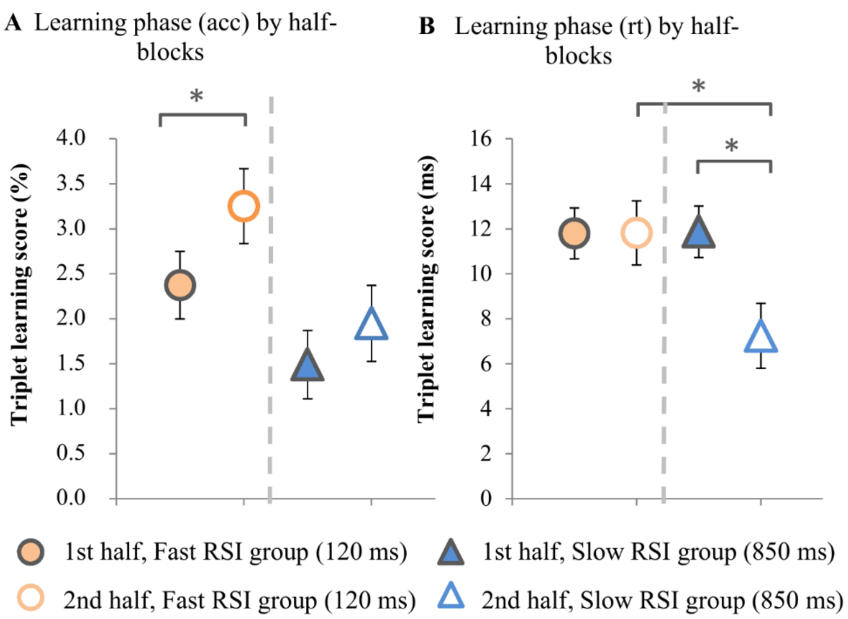


**Figure S2. The within-block position effect in the Learning phase.** The accuracy (A) and reaction time (B) triplet learning scores are presented in the Learning phase. The colors represent the groups that *learnt* with different presentation rates (orange – fast RSI group, blue – slow RSI group), and open/filled shapes represent different halves of the blocks (filled – first halves of the blocks, open – second halves of the blocks). For accuracy measures, performance in the second halves of the blocks were significantly better compared to the first halves of the blocks in the fast RSI group but not in the slow RSI group ($p$ = 0.294), although the behavioral pattern was similar in both groups and the LEARNING RSI x HALF-BLOCK interaction was not significant (see the supplementary text above). For reaction time measures, the slow RSI group showed weaker performance in the second halves of the blocks compared to the first halves of the blocks, while the fast RSI group showed similar performance in both halves of the blocks. The error bars represent SEM. ** $p < 0.01$; * $p < 0.05$; + $p < 0.1$.

**Reaction Time** – The main effect of HALF-BLOCK was significant [$F_{(1, 77)} = 6.254$, $p$ = 0.015, $\eta_p^2 = 0.075$]: the triplet learning performance was on average larger in the first halves of the blocks than in the second halves [11.831 ms vs. 9.527 ms, respectively]. The two groups' triplet learning performance differed from each other in the first and the second halves of the blocks [showed by the significant HALF-BLOCK x LEARNING RSI interaction: $F_{(1, 77)} = 6.363$, $p = 0.014$, $\eta_p^2 = 0.076$]. The LSD post hoc test revealed that the slow RSI group's triplet learning performance was smaller in the second halves of the blocks than in the first halves [$M_{\text{First half}} = 11.867$ ms vs $M_{\text{Second half}} = 7.238$ ms, $p = 0.001$; Figure

S2B]. In contrast, the fast RSI group's triplet learning performance was similar in the first and second halves of the blocks [$M_{\text{First half}}$ = 11.795 ms vs $M_{\text{Second half}}$ = 11.815 ms, $p$ =0.988]. Interestingly, the two groups' triplet learning performance was significantly different in the second halves of the blocks, with better learning performance in the fast RSI group [$p$ = 0.027]. The two groups' learning performance did not differ significantly in the first halves of the blocks [$p$ = 0.965]. The remaining interactions involving HALF-BLOCK did not reach significance [$p \geq 0.159$].

***Does the length of the RSI differentially affect performance in the first vs. second halves of the blocks in the Testing phase?***

**Accuracy** – The main effect of HALF-BLOCK and the interactions involving HALF-BLOCK did not reach significance [$p \geq 0.101$].

**Reaction time** – The ANOVA yielded a trend level HALF-BLOCK x LEARNING RSI interaction [$F_{(1, 77)} = 3.217$, $p = 0.077$, $\eta_p^2 = 0.040$], reflecting that the groups' learning scores differed in the first and the second halves of the blocks. Importantly, this effect was related to the RSI during learning but not during testing. Specifically, the post hoc tests showed that, in the first halves of the blocks, the fast RSI group exhibited smaller learning scores than the slow RSI group [7.056 ms vs. 11.846 ms, respectively, $p$ = 0.015], but there were no difference between the groups in the second halves of the blocks [9.894 ms vs. 9.987 ms, respectively, $p$ = 0.965]. The remaining main effects and interactions did not reach significance [all $p$s $\geq 0.179$].

***Summary of the within-block position effects***

Overall, the within block position effect analysis suggests that the length of RSI may have a differential effect on the expression of triplet knowledge in the first vs. second halves of the blocks, with longer RSIs leading to smaller learning scores particularly in the second halves of the blocks (at least in terms of RT data).
